# Stochastic Resonance Mediates the State-Dependent Effect of Periodic Stimulation on Cortical Alpha Oscillations

**DOI:** 10.1101/191577

**Authors:** Jérémie Lefebvre, Flavio Frohlich, Axel Hutt

## Abstract

Brain stimulation can be used to engage and modulate rhythmic activity in cortical networks. However, the outcomes have been shown to be impacted by behavioral states and endogenous brain fluctuations. To better understand how this intrinsic oscillatory activity controls the brain’s susceptibility to stimulation, we analyzed a computational model of the thalamocortical system in both the rest and task states, to identify the mechanisms by which endogenous alpha oscillations (8Hz-12Hz) are impacted by periodic stimulation. Our analysis shows that the differences between different brain states can be explained by a passage through a bifurcation combined to stochastic resonance - a mechanism whereby irregular fluctuations amplify the response of a nonlinear system to weak signals. Indeed, our findings suggest that modulating brain oscillations is best achieved in states of low endogenous rhythmic activity, and that irregular state-dependent fluctuations in thalamic inputs shape the susceptibility of cortical population to periodic stimulation.

## INTRODUCTION

Periodic brain stimulation, such as repetitive transcranial magnetic stimulation (rTMS) and transcranial alternating current stimulation (tACS), can be used to engage cortical rhythms (Frohlich 2015). Such findings have raised the fascinating prospect of manipulating rhythmic brain activity in a controlled manner, engaging neural circuits at a functional level to manipulate cognition and treat disorders of the central nervous system (Cerere et al. 2015, Frohlich 2014, Romei et al. 2016). Yet, the brain is not a passive receiver. Both invasive and non-invasive brain stimulation differ in their effect on neuronal dynamics as a function of the state of the targeted network, a robust observation across a variety of stimulation modalities (Neuling et al. 2013, Ruhnau et al. 2016, Alagapan et al. 2016). Indeed, previous studies have reported the seemingly counterintuitive fact that the susceptibility of neural tissue to exogenous control (such as during stimulation for instance) is increased during task-engaged states (Neuling et al. 2013, Ruhnau et al. 2016, Alagapan et al. 2016). Task-engaged states are generally characterized by higher firing rates and by population responses that are predominantly asynchronous, and where low-frequency electrical brain activity is suppressed compared to the rest condition (Churchland et al. 2010, Rosenbaum et al. 2016). In contrast, the rest state is characterized by strong endogenous oscillations that reflect internally driven brain processes (Pfurtscheller et al. 1996, Klimesch 2012). To compensate for these state-dependent differences in population activity, it has been suggested that stimulation parameters should be calibrated in a closed-loop (Boyle & Frohlich 2013, Lustenberger et al. 2016) in order to be optimally effective. But the lack of understanding about the cause of these state-dependent changes and how they interfere with brain stimulation remains one the main limitations to the optimization and development of new experimental and clinical paradigms based on the selective control of brain rhythms.

We formulated the hypothesis that the dominance of asynchronous activity in the task state - in the form of irregular fluctuations in neural activity - may enhance the susceptibility of cortical neurons to periodic stimulation. We notably asked whether stochastic resonance (SR) was involved. Stochastic resonance is a phenomenon by which the presence of uncorrelated random fluctuations, or so-called “noise”, amplifies the salience of a weak periodic signal driving a non-linear system (McDonnell & Abbott 2009). In absence of noise, a subthreshold signal is too weak to generate an output that reflects the input. For intermediate levels of noise, however, the superposition of the signal and the random fluctuations results in super-threshold – and thus detectable – dynamics. Further increase in noise typically causes the saliency of the signal to decrease. However, in the context of brain stimulation, it remains unclear how the task-related decrease in alpha oscillations is involved in enhancing the susceptibility of cortical neurons to entrainment by exogenous electric fields.

Computational approaches and modeling are poised to answer many of these fundamental questions, but have received little attention, especially with respect to the clinical applications of brain stimulation. By identifying the network mechanisms involved, modeling can be used to catalyze the development of more efficient stimulation protocols and customized clinical interventions. Using simulations, we have previously used a cortical network model to better understand how periodic stimulation interacts with resting-state alpha (8Hz-12Hz) activity, and found that multiple mechanisms where involved (Herrmann et al. 2016). Echoing the theory of non-linear oscillators in physics, stimulation with increasing amplitudes and/or frequencies were found to shape networks responses through different mechanisms: *resonance*, in which intrinsic oscillations are enhanced by stimuli with frequencies close to the network natural frequency; and *entrainment*, where the systems’ dynamics become phase locked to the stimulation and thus adopt the same frequency. Entrainment occurs only for specific sets of stimulation amplitudes and frequencies (regions in stimulation parameters space called Arnold tongues). But what happens in the presence of state-induced changes in baseline activity?

To address these questions and provide predictions about optimal entrainment conditions, we harnessed computational and mathematical techniques to analyze state-dependent effects on a network model of the thalamo-cortical system in presence of periodic stimulation. We found that increased thalamic drive suppressed endogenous oscillations throughout the thalamo-cortical loop, leading to an asynchronous state characterized by intense and weakly correlated spiking activity. In presence of periodic stimulation, this transition was accompanied by an increased susceptibility to entrainment, as demonstrated by amplified power at stimulation frequency and phase alignment of cortical responses with the stimulation. Specifically, we found that the oscillatory response of the system switched from alpha to stimulus-induced activity as thalamic drive was increased i.e. during a transition from resting state to the task state. Taken together, these results show that the thalamo-cortical loop implements a gain control mechanism regulating the robustness of alpha oscillatory activity and by doing so, modulates cortical susceptibility to rhythmic entrainment by brain stimulation.

**Figure 1.**
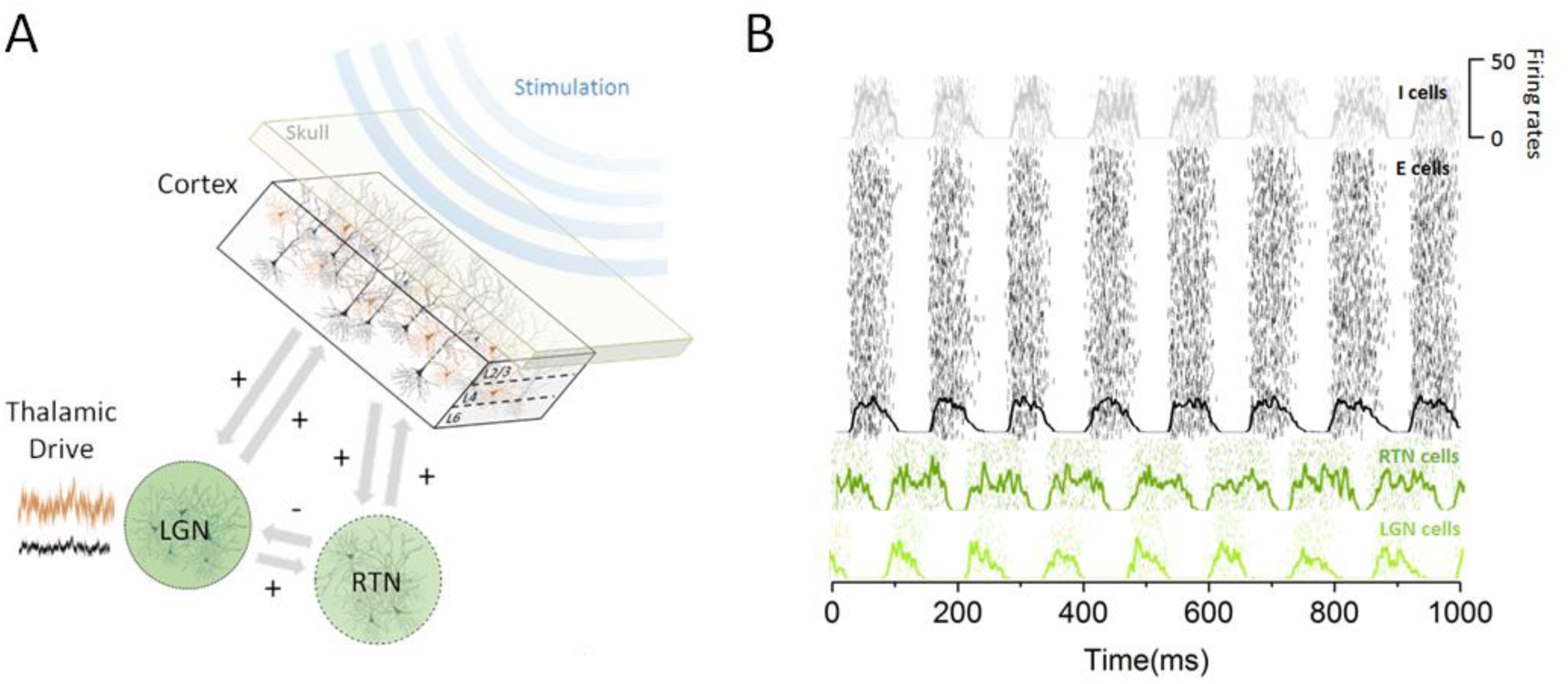
Thalamo-cortical circuit schematic and resting state activity. **A.** Our computational model of the thalamo-cortical system is composed of both cortical (excitatory and inhibitory) and sub-cortical (lateral geniculate nucleus (LGN) and reticular(RTN)) populations. Sensory inputs, whose intensity scales with state, is assumed to arrive at the thalamus to then influence the dynamics of the whole circuit. Non-invasive periodic stimulation is applied only to cortical neurons. **B.** In the resting state, both cortical and sub-cortical neurons are maintained in a deeply synchronous regime, displaying phase locked firing rate oscillations in the alpha band (8Hz).

## RESULTS

To understand how the efficacy of periodic stimulation could depend on brain state (Alagappan et al. 2016) we examined the hypothesis that cortical susceptibility to entrainment is controlled by sub-cortical (i.e. thalamic) populations. Given the important role of the thalamus on cortical state control (Poulet et al. 2012) and resting state alpha activity (Hughes et al. 2004, Hughes et al. 2005, Lorincz et al. 2009), we investigated how increased thalamic inputs regulated cortical excitability and the power of endogenous alpha oscillations. We thus developed a model of the thalamo-cortical network composed of cortical, thalamic relay and reticular populations that exhibit synchronous activity within the alpha band in the resting state (see Materials & Methods). In this model, cortical populations of excitatory and inhibitory cells share both feedforward and feedback connections with the thalamus. Visual inputs (or other sensory inputs meant to fluctuate as a function of state) are sent through the LGN to cortical areas for further processing (Poulet et al., 2012), impacting the pattern of activity of cortical neurons. The model structure is illustrated in Figure 1, along steady state responses generated by the different groups of neurons present.

**Figure 2.**
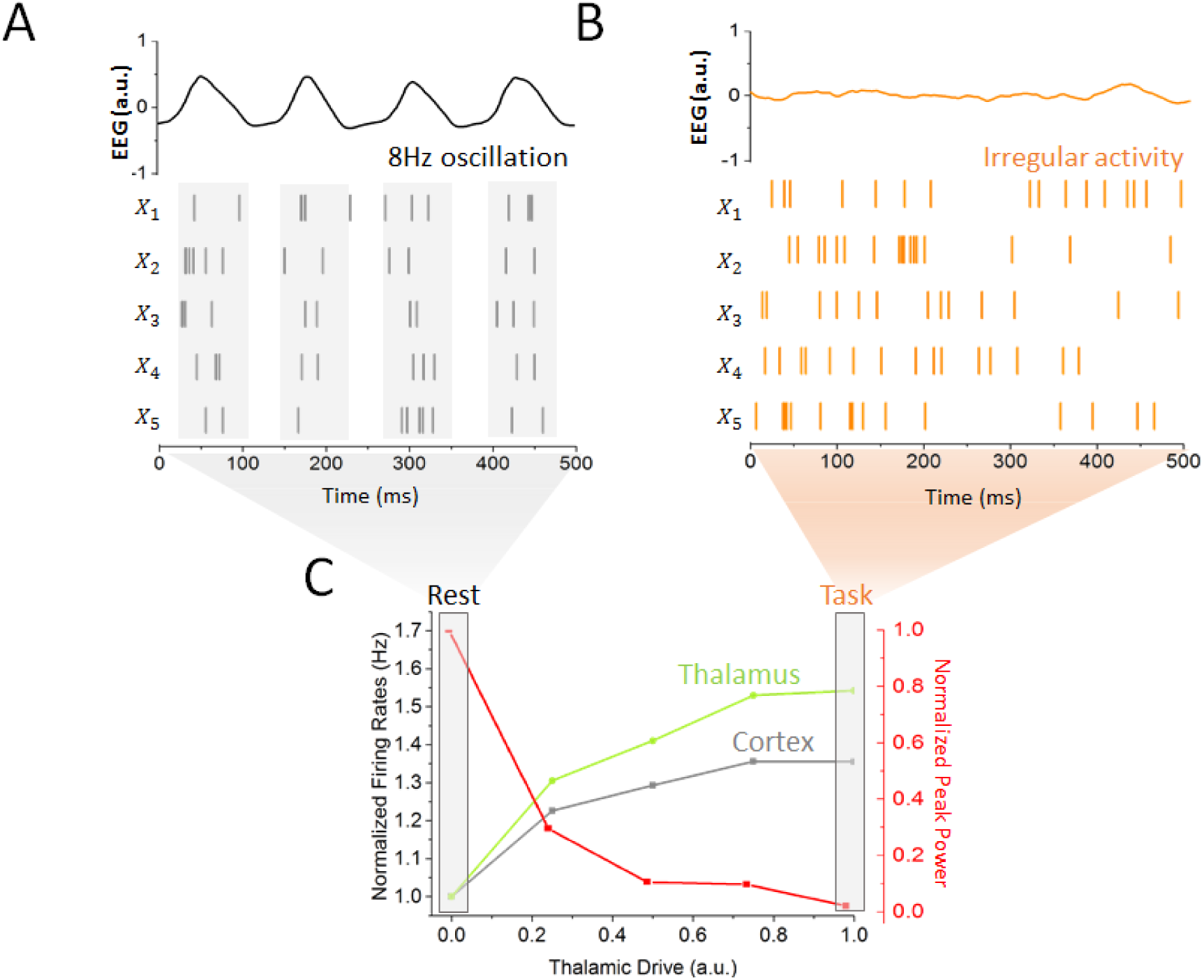
Impact of thalamic drive on firing rates and alpha power without stimulation. **A.** Sample spiking activity of 5 randomly selected excitatory cortical cells in the resting state, which is defined as a regime of minimal thalamic drive (*D*_*LGN*_ *=* 1*x*10^-4^). The firing activity of those neurons is highly correlated, and synchronized to the phase of simulated EEG alpha oscillations (top). **B.** Activity of the same cortical neurons in the task state, where thalamic drive is high (*D*_*LGN*_ *=* 1). The spiking activity is now irregular and asynchronous, leading to a flat EEG activity where alpha oscillations have been suppressed. C. Effect of gradual increase in thalamic drive on alpha power and mean firing rates for both cortical (grey) and sub-cortical (green) neurons.

To first quantify the impact of thalamic drive on cortical activity and resting state alpha oscillations - first without stimulation - we measured both cortical and sub-cortical firing rates and the power of endogenous oscillations as input to the thalamus was increased (Figure 2). For minimal inputs to the thalamus, all populations across the network were maintained in a highly synchronous state, where recurrent interactions generate strong firing rate oscillations within the alpha band, with an envelope of about 8Hz. As can be seen in Figure 1A, the activity of the different populations was phase locked, where the phase difference was due to the presence of propagation delays from the thalamus to and back from the cortex. However, increases in thalamic inputs had a destabilizing effect on endogenous oscillations. While increasing thalamic drive also increased firing rates throughout the system, a gradual suppression of synchronous alpha activity could also be observed: the power of endogenous alpha oscillations decreased significantly. This increased thalamic drive also significantly changed the spiking patterns of cortical neurons: spiking activity became irregular and asynchronous, in line with studies showing a transition towards decorrelated cortical dynamics at task-onset (Pfurtscheller & Lopes da silva 1999, Poulet et al. 2012, Churchland et al. 2010). Taken together, these results demonstrate that in our model, changes in thalamic state is sufficient to destabilize resting state alpha oscillations throughout the thalamo-cortical loop as well as shape spike correlations. To better distinguish these two cases, we defined the rest and task states as limit cases of low (i.e. rest) and high (i.e. task) thalamic drive, respectively (see Materials & Methods), as indicated in Figure 2C. We note that aside from the thalamic input which was changed to set the system in either the rest and/or task states, only stimulation parameters, such as amplitude and frequency, were changed. All other network parameters were kept constant. Such a smooth and gradual destabilization of alpha oscillations is suggestive of a noise-induced supercritical Hopf bifurcation (See Material and Methods). According to this well-known mechanism (Horstenke & Lefever), noise acts as a gain control parameter by which non-linear oscillations sustained by delayed and recurrent inhibition (as in our model) can be altered and/or suppressed by a change in noise variance (Lefebvre et al. 2015, Hutt et al. 2016). In the process, noise also linearizes the dynamics by smoothing the non-linear response function of threshold systems (Gammaitoni 1995, McDonnell & Abbott 2009, Lefebvre et al. 2015). In the context of our model, rest and task states would then correspond to opposite points above and below the transition point, where the “noise” is here mediated by increasingly irregular input from thalamus to cortex.

**Figure 3.**
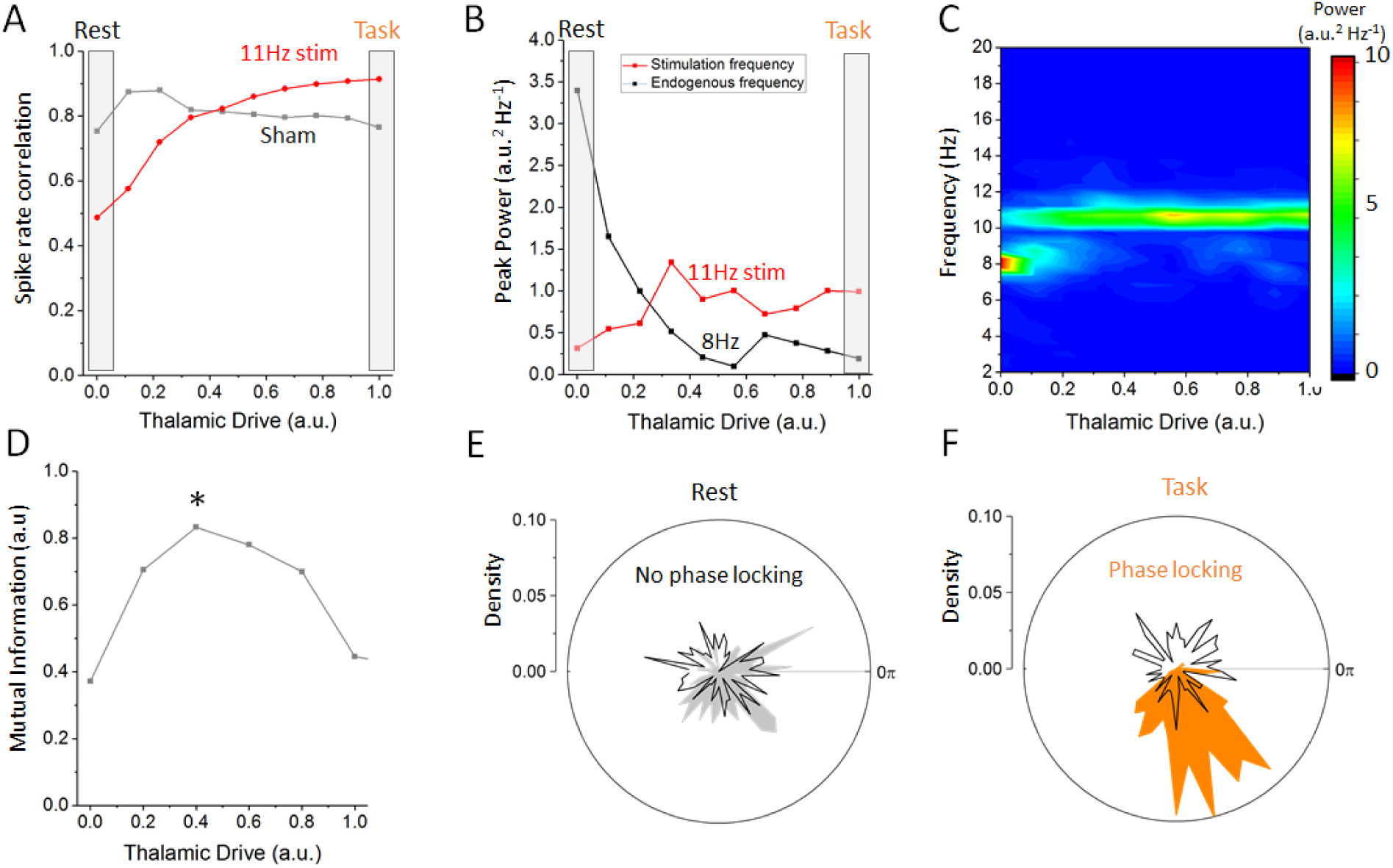
Effect of increasing thalamic drive on network dynamics with stimulation. **A.** Firing rate correlations of cortical excitatory neurons (average pairwise correlation between firing rates) as a function of thalamic drive. Correlations increase monotonically with thalamic input as the network gets gradually entrained by the stimulation. B. Peak power measured at the endogenous alpha frequency (*ω*_0_=8Hz) and at the stimulation frequency (*ω*_*stim*_=11Hz) as a function of increasing thalamic drive. Power of endogenous oscillations is gradually suppressed, while the opposite occurs at the stimulation frequency, suggesting a transition between oscillatory regimes. Note the presence of a peak in the power at the stimulation frequency, suggesting an optimal value of thalamic drive at which cortical activity gets entrained. C. Full power spectral density distribution as a function of thalamic drive. Following the destabilization of endogenous oscillations, spectral power only remains at the stimulation frequency. Stimulation parameters for A, B and C were *S =*0.15, *ω_stim_*=11Hz. D. Mutual information between stimulation and network response as a function of increasing noise to the thalamic population. A stimulation waveform of weak amplitude (here *S =*0.05) is applied, and the mutual information displays a maximum at some intermediate value of noise – indicating that stochastic resonance is involved. E. Distribution of phases across trials of cortical firing rates across trials at the stimulation frequency (11Hz) in the rest state (*D*_*LGN*_ *=* 1*x*10^-5^). Stimulation is applied at a random, uniformly distributed phase across trials (black line). The averaged phase difference between stimulation and cortical firing rate is uniform, indicating that the network dynamics are not phase locked to the stimulus. F. In contrast, in the task state (*D*_*LGN*_ *=* 1*x*10^-5^), cortical firing rates are phase locked to the stimulation and the distribution shows a peak at preferred phase (orange area). Here *S =* 0.1 and *ω*_*stim*_=11Hz.

Having defined rest and task states in our network model and the influence of thalamic activity on alpha oscillations, we next asked how state-dependent changes in resting state oscillations (the destabilization of alpha oscillations) were impacted by cortical periodic stimulation. To disambiguate the contribution of resonance (the amplification of endogenous oscillations) and entrainment (the phase locking to an externally applied frequency) and test network responses, we first considered a cortical stimulation with moderate amplitude and frequency of of 11Hz (*ω*_*stim*_), which does not share any low order harmonic relationship with the endogenous frequency (i.e. *ω*_*o*_ *=*8Hz). This was done to align our simulation our previous experimental setting (Alagapan et al. 2016), to see if we could reproduce state dependent effect observed. We then computed how spiking activity and cortical oscillations properties changed as thalamic drive was gradually increased in presence of periodic stimulation (Figure 3). We found that cortical spike rate correlations – induced by this stimulation – scaled with the input to the thalamus. Compared to the non-stimulated case, spike rate correlations decreased in the rest state, indicating a desynchronization of cortical activity. In contrast, as thalamic drive was gradually increased towards the task state, spike rate, correlations increased again, suggesting that cortical neurons are getting synchronized, but this time to the stimulation and not to an internal attractor (in this alpha oscillations). These observations indicate that there is a gradual transition from recurrent to externally driven dynamics as thalamic input is increased. To explore this further, we measured in Figure 3B the peak power found both at alpha frequency (8Hz) and stimulation frequency (here 11Hz). We found that the power measured at stimulation frequency indeed increased during the transition from the rest to task state and that a shift in oscillatory regime can be observed. Indeed, one can clearly see a jump in dominant frequency from 8Hz (endogenous alpha frequency) to 11Hz (stimulation frequency) as resting state alpha oscillations are gradually suppressed.

Interestingly, we found a value of input for which the peak power at stimulation frequency is maximal, suggesting that enhanced spectral power at the stimulation frequency might occur in our model through stochastic resonance. Stochastic resonance (SR) has been implicated in the enhancement of sub-threshold signals in which random fluctuations – of either external or internal origin - act like a pedestal increasing the sensitivity of stimulated neurons to a given set of low intensity inputs (Miniussi et al. 2013). Initially formulated in bistable systems with an implicit time-scale (a “resonance”), SR is also present in systems with threshold nonlinearities (McDonnel), which we note is present in our model.

To verify whether SR was indeed involved – and disambiguate the contribution of SR and dithering (McDonnell & Abbott 2009) - we chose a weak amplitude stimulation and computed the mutual information between stimulation signal and firing rates response at the stimulation frequency across successive and independent trials. As shown in Figure 3D, the mutual information was found to be optimized at an intermediate value of noise, indicating that SR - in its most strict definition (McDonnell & Abbott 2009) - was indeed involved. To confirm the presence of entrainment (to the stimulation and not to another internal oscillation), we performed a phase analysis comparing the phase of the cortical firing rates with the phase of the stimulation across independent trials. The results are shown in Figure 3E and 3F. There, one can see that weak phase relationships were found in the rest state (3E) while clear phase locking can be seen in the task state (3F). Our simulations show that cortical spiking responses are phase locked to the stimulation waveform only for intermediate values of thalamic drive. Below this drive, endogenous network activity suppresses cortical entrainment, and beyond this value, irregular noisy fluctuations originating from the thalamic population decreases the saliency of the evoked responses. Taken together, the results above indicate that state-dependent inputs simultaneously amplify stimulation-induced activity while suppressing endogenous rhythmic activity through stochastic resonance, enabling the stimulation to entrain cortical neurons.

**Figure 4.**
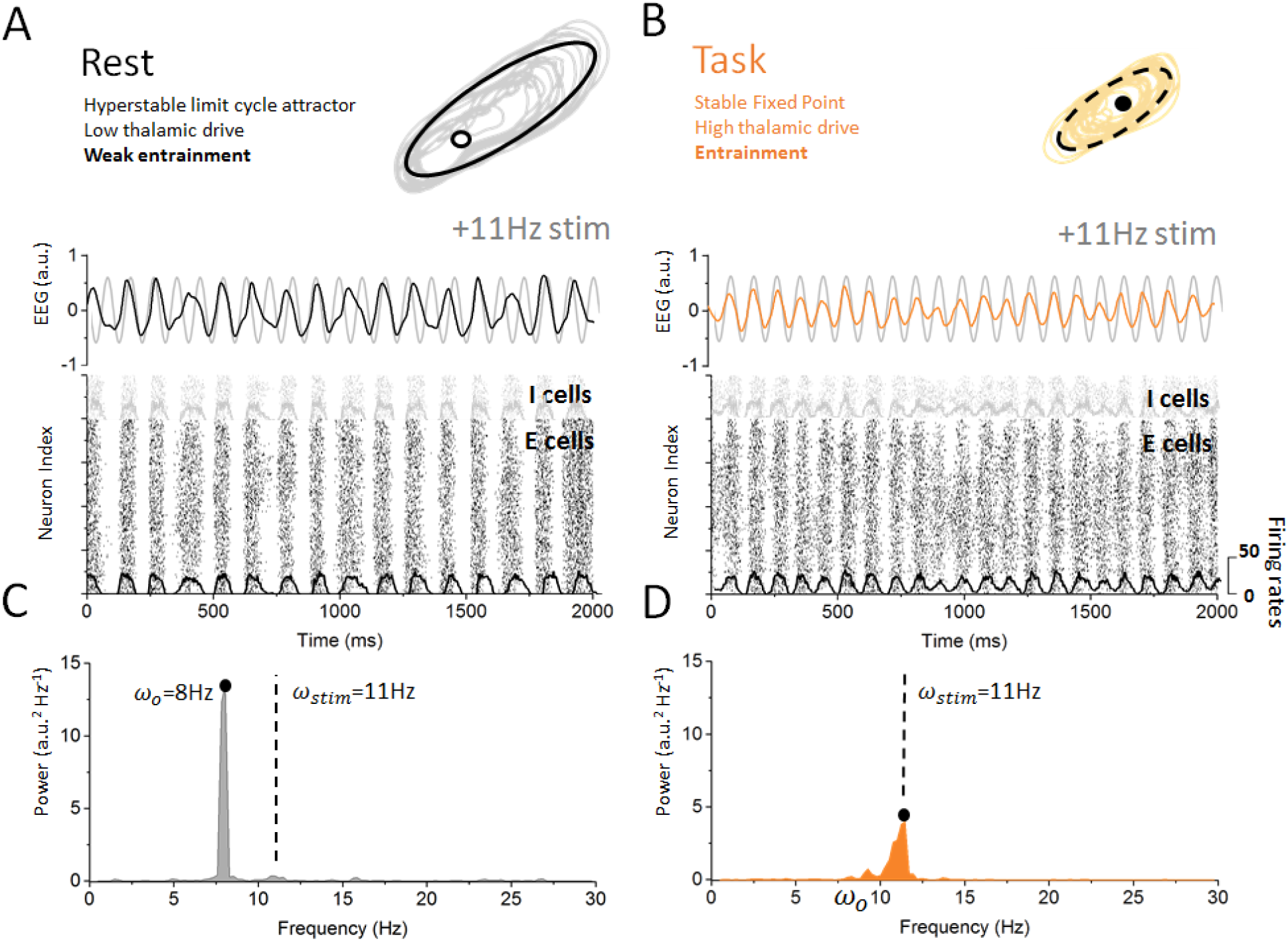
Response of cortical neurons to periodic stimulation in the rest and task states. A. The resting state is characterized with strong synchronous alpha activity, leading to stable attractor dynamics and resulting in weak entrainment of endogenous oscillations. Despite the presence of periodic stimulation impacting equally all cortical neurons, the dynamics of the simulated EEG is predominantly characterized by endogenous oscillations (top). The spiking activity of both excitatory and inhibitory neurons remains locked to endogenous cycles where stimulation has little to no impact on network activity (bottom). **B.** In contrast, endogenous oscillations are suppressed in the task state where the dynamics evolve around a fixed point. Simulated EEG activity is fully entrained to the stimulation (top)and so are cortical neurons whose spiking is phased locked to the stimulation frequency**. C.** Power spectral density distribution of the EEG response in the rest state. The spectrum is largely dominated by endogenous oscillations (*ω*_*0*_ *=*8Hz) and only weak contribution can be observed at the stimulation frequency (*ω*_*stim*_ *=*11Hz), indicating that network oscillations are not entrained by the stimulation. D. Power spectral density distribution of the EEG response in the task state. Here, in contrast, the power at the endogenous frequency has been almost fully suppressed and a clear peak can be seen at the stimulation frequency. This implies that network oscillatory activity is fully determined by the stimulation. Stimulation parameters are here *S =*0.15, *ω*_*stim*_=11Hz.

Figure 4 summarizes these findings in the rest and task states. In the rest state (Figure 4A), the dynamics of the cortical neurons were predominantly governed by internal, recurrent fluctuations, as the stimulation did not significantly entrain the alpha oscillations. In this state, applied stimulation has minimal impact on the activity of cortical neurons: the phasic alternation of network-wide coincident firing and inhibitory recovery greatly reduces susceptibility to entrainment. This absence of phase locking of neuronal activity to the stimulation signal can be most clearly seen from the firing activity of cortical excitatory and inhibitory neurons. The firing activity of these cells occurs by successive bursts at 8Hz, with little effect from stimulation-induced membrane fluctuations, which remain sub-threshold. The poor susceptibility is also apparent from the relatively small contribution of stimulation power to the overall spectral density distribution of cortical EEG (Figure 3C). Taken together, these simulations indicate that the rest state is characterized by highly stable attractor dynamics. In contrast, robust entrainment was observed in the task state (Figure 4B), as seen from a dominant spectral peak located precisely at the stimulation frequency (Figure 3D). Despite the fact that stimulation amplitude and frequency remained the same, cortical neurons were here fully entrained by the 11Hz stimulation. The irregular activity that characterizes the task state was here replaced by stimulus-induced fluctuations while neural firing was tightly phased locked to the stimulus dynamics. Spectral power at the stimulation frequency is significantly amplified compared to the rest state, while negligible power can be found at the endogenous alpha frequency. This forms yet another indication that the resting state attractor has been destabilized under the action of thalamic drive and resulting increase in neural noise (Faisal, Lefebvre et al. 2015, Hutt et al. 2016), also in line with resonance (McDonnell & Abbott 2009).

**Figure 5.**
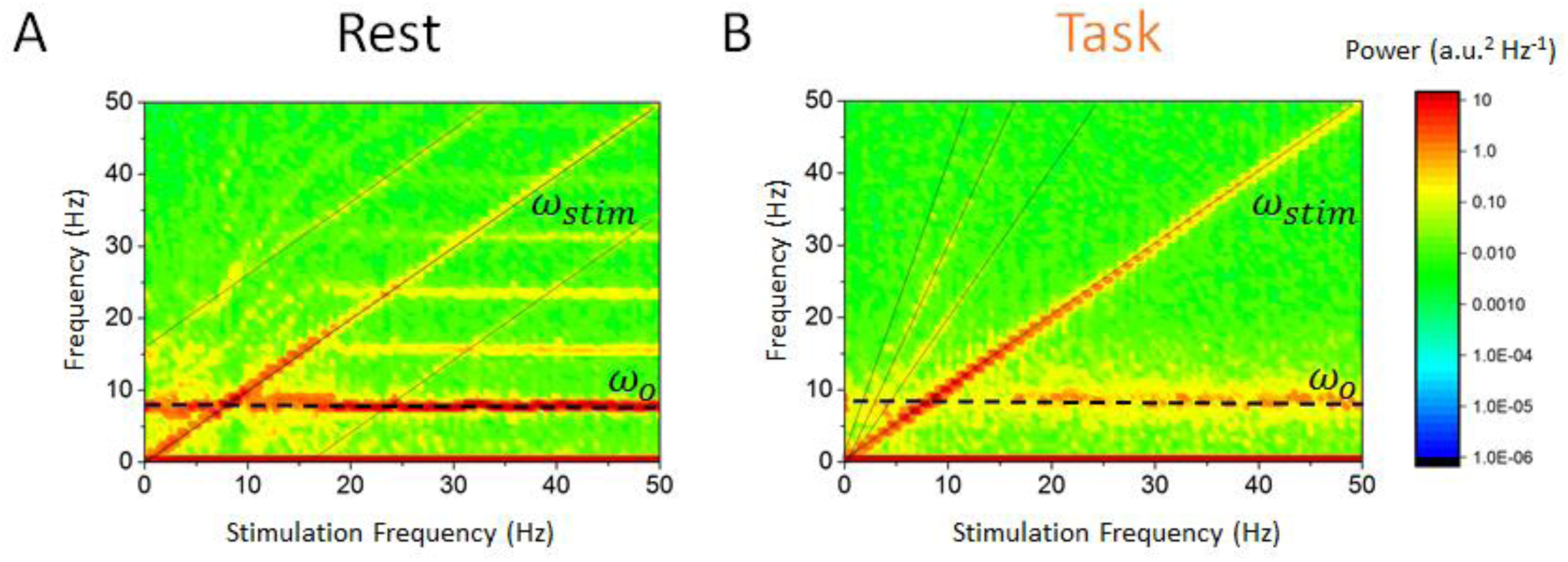
State-dependent responses to periodic stimulation with variable frequency. **A**. Power in the rest state is concentrated at the endogenous frequency (horizontal dashed line at 8Hz) while the stimulation frequency *ω*_*stim*_ is increased from 0 to 50Hz. The peak power found at the stimulation frequency (dashed line along the diagonal) is small unless *ω*_*0*_ ≈ *ω*_*stim*_, that is, when the stimulation frequency is close to the peak alpha frequency. **B.** In the task state, power is instead concentrated at the stimulation frequency across the range of values visited (along diagonal). In contrast, power at the endogenous oscillation is much smaller. Note that the power at the stimulation frequency scales with the distance with respect to the alpha peak at 8Hz. Stimulation parameters are here *S =*0.15 while *ω*_*stim*_ was varied from 0 to 50Hz.

But how does this state-dependent susceptibility depends on stimulation parameters? To answer this question, we first fixed stimulation amplitude and varied its frequency between 0 and 50Hz while measuring the power spectral density in each case (Figure 5). While a peak at the stimulation frequency can be seen along the diagonal, the rest state power spectral density is characterized by a dominant horizontal peak at 8Hz. According to this analysis, the spectral contribution of stimulation is negligible unless stimulation frequency is close to the endogenous frequency and/or its harmonics (horizontal lines in Figure 5A), as expected via resonance (Ali et al. 2013, Herrmann et al. 2016). In the task state, the opposite occurs: reduced power at the endogenous frequency promotes entrainment and stimulation power increases significantly across the range of stimulation frequency considered.

**Figure 6.**
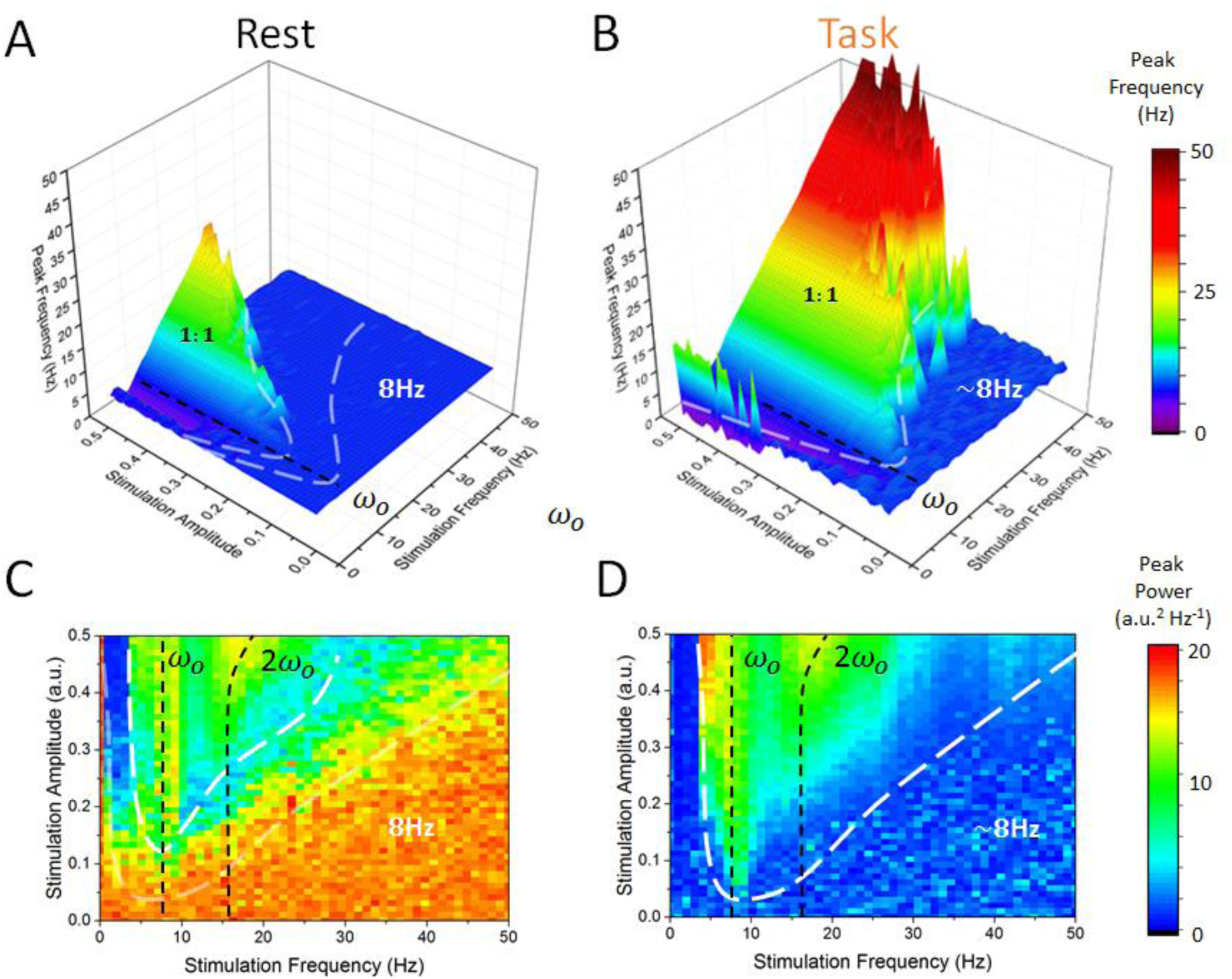
State-dependent Arnold tongues as a function of stimulation amplitude and frequency. By systematically varying the stimulation amplitude (*S*) and frequency (*ω*_*stim*_) and measuring peak frequency (i.e. frequency where maximal power can be found) and peak power (i‥e power at the peak frequency) in each case, we can delimit regions in parameter space where cortical dynamics is governed either by endogenous alpha oscillations or locked to the stimulation. **A.** In the rest state, the vast majority of stimulation parameter space is characterized by an absence of entrainment. The peak frequency of cortical EEG activity remains at 8Hz. However, as stimulation amplitude in increased for stimulation frequency near 8Hz, entrainment occurs. This triangular region – called the Arnold tongue – remains however narrow. **B.** In the task state, the Arnold tongue increases significantly, and occupies most of the range of stimulation parameters considered – entrainment is thus more prevalent in the task state. **C.** Peak power measured in the rest state for varying stimulation parameters. **D.** Same as in C for the task state. Stimulation amplitude was varied within [0, 0.5] and its frequency within [0, 50Hz].

As a next step, we computed the power-spectral density of the simulated EEG activity of cortical excitatory and inhibitory cells and tsystematically measured the peak frequency and power for all combination of stimulation frequency and amplitudes within a given range. We then identified regions for which the dominant (i.e. peak) frequency was defined either by endogenous oscillations or by the stimulation – thus identifying regimes of entrainment. In Figure 6A, one can see the peak frequency of the simulated cortical EEG activity in the rest state. For most combinations of stimulation amplitudes and frequencies, the peak frequency remains stable and equal to the endogenous alpha frequency (i.e. 8Hz). However, for higher amplitudes and for stimulation frequencies near 8Hz, cortical neurons are gradually entrained by the stimulation (1:1 entrainment). A narrow triangular entrainment region (the so-called Arnold tongue, delimited by white dashed lines) (Glass & Belair 1980, Hunter & Milton 2003) emerges and gets larger as stimulation amplitude increases. Note that the asymmetrical shape of the Arnold tongue is here due to a stimulation-induced shift of the endogenous frequency (Hutt et al. 2016). The equivalent calculations were made in Figure 6B but for the task state. Multiple differences compared to the rest state can be readily be seen. The Arnold tongue spans a much larger portion of stimulus parameter space and indicates robust entrainment for much lower values of stimulation amplitudes. Outside the Arnold tongue, where there is no entrainment, the increase in thalamic input yields more noisy and irregular dynamics, leading to shifts in the estimates for the endogenous frequency (Lefebvre et al. 2015). In Figure 6C and 6D, the power at peak response frequency is plotted. The power found at the endogenous frequency in the rest state outside the Arnold tongue (here also delimited by a white dashed line) is high, and is gradually suppressed as stimulation parameters are changed, towards the entrainment region. There, peak power, associated with the stimulation frequency is much smaller (Figure 6C). Under the action of thalamic input in the task state, endogenous alpha oscillations are suppressed, leading to a weak power outside the Arnold tongue (Figure 6D). In both task and rest states, peak power is increased due to resonance if the stimulation frequency is near the endogenous frequency (vertical black dashed lines in Figures 6C and 6D). Note here also the slight shift in resonance frequency for higher stimulation amplitudes (Lefebvre et al. 2015).

But what mechanism is responsible of this state-dependence? Mathematical insights can play here a key role in understanding how periodic stimulation interacts with intrinsic limit cycles (i.e. alpha oscillations) and how changes in stability can promote entrainment. In (Alagapan et al. 2016), the authors modelled state-dependent entrainment using a population-scale network of cortical neurons interacting through both local and feedback projections and with robust resting state oscillations. To obtain some more insight about the mechanism involved, we there considered a simplified neural oscillator with delayed feedback (Lefebvre et al. 2013, Lefebvre et al. 2015, Hutt et al. 2016) as simplified model for the thalamo-cortical network we previously considered. Networks of neurons exposed to delayed feedback and recurrent inhibition commonly display rhythmic activity whose features are tightly linked to input statistics (Dhamala et al. 2001, Doiron et al. 2003, Lefebvre et al. 2013). To better understand the mechanism underlying the state-dependent changes in the amplitude of resting state oscillations, we investigated the dynamics of this simplified model (see Materials & Methods) in presence of periodic forcing (i.e. stimulation) and various levels of noise. In can be shown that as input noise increases during the task state, limit cycle solutions are destabilized through a supercritical Hopf bifurcation. Analyzing the response of this simplified model both above and below the bifurcation threshold, that is, in the vicinity of the point where self-sustained (i.e. intrinsic) oscillations become unstable, we found that limit cycle solutions of this system are entrained by periodic forcing although their amplitudes are highly sensitive on noise intensity. To see this, we computed in Figure 7B resonance curves for the equation and compared the results for both high and low values of input noise variance. This simple qualitative analysis shows that the linearization due to noise decreases the amplitude of the resonant solutions and increases the amplitude of all other forcing frequencies by transitioning through a critical state. This spectral clustering in which power and/ or amplitude of forced solutions is concentrated near the resonance is a state of weak susceptibility to entrainment. The consequences of input-induced linearization in the simple conceptual model above is two-fold: 1) it causes a suppression of resonant oscillations, and 2) augments the amplitude of non-resonant solutions. This was found to be in agreement with the state-dependency observed in our initial model as depicted in Figure 7C.

**Figure 7.**
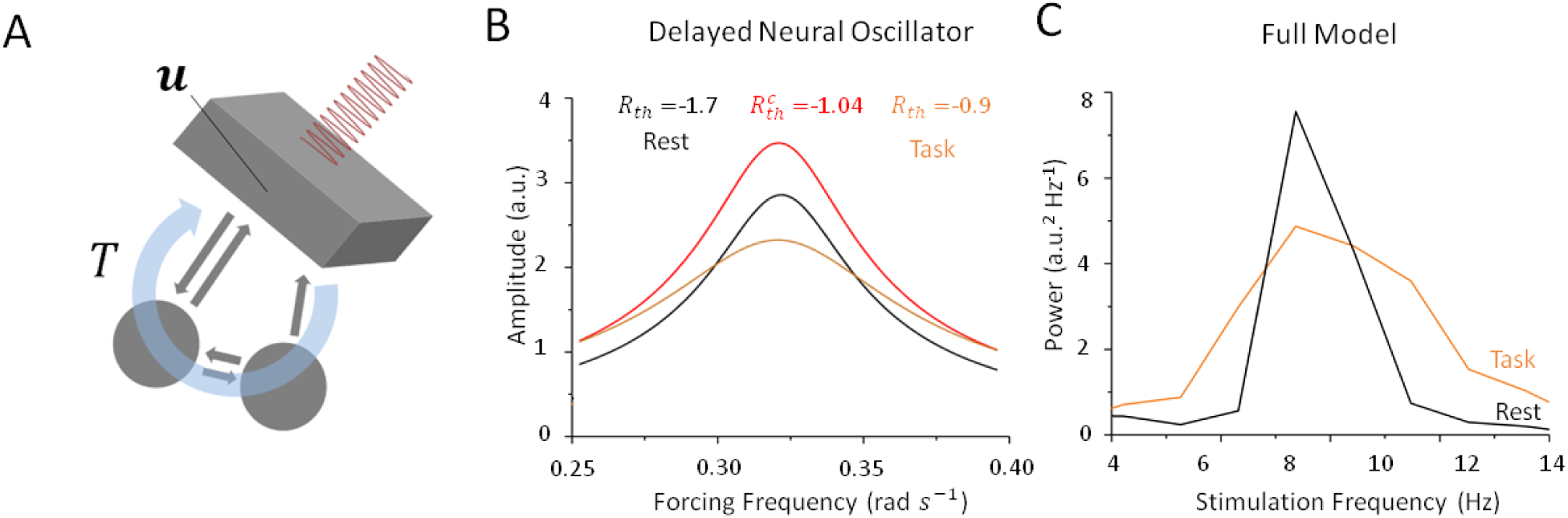
Stochastic resonance in a noisy and delayed neural oscillator. **A.** We investigated the dynamics of a conceptual delayed feedback model in presence of periodic forcing (i.e. stimulation) as a proxy for the thalamo-cortical circuit subjected to additive noise. The mean activity *u* experiences delayed feedback with delay *T*, additive noise and periodic forcing. **B.** Amplitude of entrained solution both above and below the Hopf bifurcation threshold. Below the bifurcation (black line), the linear gain |*R*_*th*_| remains high because noise is intensity is small. Solutions have a high amplitude only near the intrinsic critical frequency. Above the bifurcation (orange line), the linear gain |*R*_*th*_| becomes smaller under the effect of noise. As a consequence, the amplitude of all non-resonant solutions increases, while the amplitude of resonant solution decreases. **C.** Peak power at *ω*_*stim*_ in the rest (black) and task (orange) states as stimulation frequency varies in the vicinity of *ω*_0_. While resulting from a more complex model, the effect remains qualitatively similar to the simplified case in B.

## DISCUSSION

Alpha activity has been implicated in a wide variety of physiological and cognitive functions (Basar 2012, Mierau et al. 2017). Numerous recent studies have demonstrated that periodically modulated electric fields can interfere with alpha oscillations to trigger measurable effect on stimulus perception (Chanes et al. 2013, Cecere et al. 2015) and task performance by entraining these rhythms (Henry et al. 2014, Samaha & Postle 2015). However, the efficacy of these approaches have been shown to depend heavily on brain states and fluctuates according to internally governed dynamics (Zaehle 2013, Minuissi et al. 2013, Ruhnau et al. 2016, Alagapan et al. 2016).

To better understand this phenomenon and elucidate previous experimental findings (Alagapan et al. 2016), we here combined computational and mathematical approached and analyzed a computational model of the thalamo-cortical system exhibiting resting state alpha oscillations. The dynamics of this network model was scrutinized in two conditions: the rest state – which we have defined as a regime of weak inputs to the thalamus, and the task state – where thalamic drive is high. We found that a destabilization of alpha oscillations occurred during the transition between rest and task states, in which both cortical and sub-cortical spiking patterns switched from correlated and synchronous to irregular and asynchronous. Then, applying periodic stimulation to entrain cortical neurons, we found that the susceptibility of the system to external perturbations was strongly controlled the presence (or absence) of an oscillatory attractor (i.e. the alpha oscillation) and as such depended on state. By comparing the stimulation waveform with the cortical firing rate responses, for weak simulation amplitudes using mutual information, we found that stochastic resonance (SR) was involved in improving cortical susceptibility to entrainment. Indeed, while in the rest state, applied stimulation had minimal impact on the activity of cortical neurons whose dynamics remains locked to internal, recurrent oscillations. Arnold tongues (i.e. regions in stimulation parameter space where network activity is phase locked to the stimulation) were found to be very narrow. In contrast, in the task state, excitatory and inhibitory cortical neurons were freed from the intrinsic attractor and found to be more susceptible to entrainment, as shown by significantly larger Arnold tongues. Using a simplified network model, our mathematical analysis revealed that thalamic input to the cortex, playing a similar role as noise, could control the stability of intrinsic oscillations by provoking a bifurcation, and by doing so, amplify the response to non-resonant stimuli. Aside from the state-dependent input and/or noise, only stimulation parameters, such as amplitude and frequency, were changed in our study. Taken together, our results show that internal control over attractor states shapes the susceptibility of cortical populations to stimulation and entrainment. Our modelling results suggest that the thalamo-cortical system behaves like a non-linear oscillator whose susceptibility to periodic forcing is gated by noise through stochastic resonance and a passage through a bifurcation.

Stochastic resonance is widely appreciated as a mechanism that regulates cortical excitability (Miniussi et al. 2013) and has measurable effects on cognitive performance and perception (van der Groen & Wenderoth 2016). Our computational results demonstrate the important role of SR, and thus of asynchronous and irregular neural dynamics, in regulating responses of cortical neurons to stimulation. Using a simplified neural oscillator model, we’ve shown that noise – playing the role of uncorrelated inputs from the thalamus – can suppress oscillation through a Hopf bifurcation. Practically speaking, this is accomplished by a noise-induced linearization of the system’s response function, a generic effect observed in systems with threshold non-linearities (see McDonnell & Abbott 2009 and references therein), which can further be shown to impact the frequency and amplitude of oscillatory solutions in non-linear systems (Lefebvre et al. 2015, Hutt et al. 2016). Such linearization is also playing a fundamental role in dithering, a mechanism exploiting SR by which noise is added to a signal, prior to its digitalization, to increase its integrity (McDonnell & Abbott 2009). Formulated in the context of neural systems, irregular and/or uncorrelated fluctuations can thus be used to increase the saliency or fidelity of neural responses. Interestingly, this “correlation-induced blockade” has been previously discussed with respect to alpha oscillations (Klimesch et al. 2007), where large-scale synchrony has been hypothesized to implement a functional inhibition gate through which increased synchronized fluctuations shuts down the mutual coding capacity of neural networks (Zohary et al. 1993). In this perspective, our results suggest that alpha oscillations – as well as other intrinsic attractors - block perturbations of neural activity, triggered either by endogenous (e.g. inter-area signaling) or exogenous (e.g. stimulation) manipulations.

In summary, our results demonstrate that target engagement by stimulation is state-dependent. The fact that the presence of a pronounced endogenous oscillation limits the impact of stimulation in terms of entrainment points towards important questions for the use of such stimulation paradigms in research, and ultimately in the clinic. First, the overall state of the network needs to be measured and documented in studies that investigate brain stimulation for the modulation of brain rhythms. Second, it remains mostly unclear how adjusting the frequency of the alpha oscillation affects cognition and behavior (Romei et al. 2016, Mierau et al. 2017). Yet, a reduced alpha peak frequency is a hallmark of a range of neurological and psychiatric illnesses (e.g. Rossini et al. 2007) and likely an overall marker of brain health (Klimesch 1999, Klimesch 2012, Vlahou et al. 2014). In the future, stimulation paradigms may be designed to alter to increase the peak frequency. In such work, our findings suggest that states of low alpha oscillations represent the ideal state for the application of stimulation. Overall, our findings suggest that modulating brain oscillations is best achieved in states of low endogenous rhythmic activity, a finding that may be important for the next generation of brain stimulation paradigms developed using rational design.

## MATERIALS AND METHODS

### Model

We have developed and analyzed a model of the thalamo-cortical system in which recurrent interactions between the different neural populations generate strong synchronous activity within the alpha band (8-12Hz). An illustration of the model structure and connections is presented in Figure 1A. The cortical population is composed of recurrently connected excitatory pyramidal neurons and inhibitory interneurons, and both interact with thalamic and reticular populations via delayed connections. The spiking activity of all neurons is modeled by a non-homogenous Poisson process,

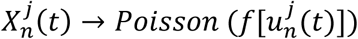

where 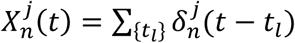 is the spike trains of the *j*^*th*^ neuron in the population *n =*{*e*, *i*, LGN, RTN}. The activation function *f*[*u*] *= f*_0_(1 + exp{-β(*u* - *h*)})^-1^ represents a saturating firing rate function that relates monotonically to the cellular membrane potential *u*, with gain *β*, maximal rate *f*_*0*_ and threshold *h*. The mean somatic membrane potentials 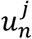 of all neurons in the network obey the set of dynamic equations

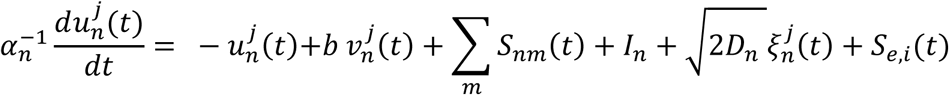

where we have spike frequency adaptation 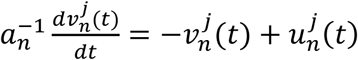 and where recurrent inputs are

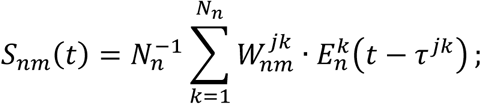

The rates *α*_*n*_ and *a*_*n*_ define the time scale of the somatic membranes and adaptation, respectively. The post-synaptic potentials 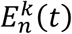 are computed by convoluting spike trains with exponential synapses with time constant *τ*_m_ i.e.

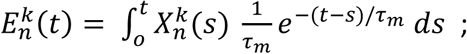

All populations, including the sub-cortical neurons (relay, reticular), are mutually coupled with sparse and spatially topographic projections (Hellwig 2000). The synaptic connectivity kernels between neurons of index *j* and *k* respectively from population m and *n* are given by Gaussians

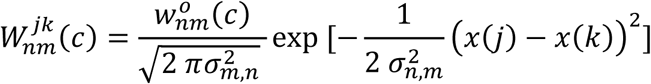

with connexion probability *c =* 0.2 and 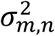 are the spatial spread of the niterneuronal connections‥ The network is also subjected to propagation delays *τ*^*jk*^ *=* |*x*(*k*) - *x*(*j*)| *v*^-1^ due to finite axonal conduction velocity *v*=0.35m/s. Furthermore, a delay of *τ*_*th*_ *=*45ms was included between thalamus (RTN, LGN) and cortex (*e*, *i*), and a delay of *τ*_*rtn*_=10ms between reticular (RTN) and relay (LGN) populations.

### Inputs and state-dependent thalamic drive

In addition to fixed bias inputs *I*_*n*_, neurons in the network are subjected to Gaussian white noise *ξ*_*n*_ of intensity *D*_*n*_. To represent an increase in sensory afference to the thalamus, the intensity of the noise driving the lateral geniculate nucleus (LGN) neurons is increased. We analyzed the dynamics of this network model in two conditions: the rest state – which we define as a regime of low thalamic drive (i.e. *D*_*LGN*_ *=* 1 × 10^-4^), and the task state – which is defined as regime of high thalamic drive (i.e. *D*_*LGN*_ *=* 1). Specifically, a transition between the rest and task states occurs whenever noise intensity at the LGN increases. Aside from this input which was changed to set the system in either the rest and/or task state, only the periodic stimulation parameters, such as amplitude and frequency, were changed. We chose not to define the thalamic drive intensity for the task state as the optimal value found through stochastic resonance (i.e. Figure 3B and 3D), because this value was found to be sensitive on the stimulation amplitude and frequency, which will vary in the subsequent analysis. To represent the effect of periodic stimulation, all excitatory and inhibitory cortical neurons were equally driven by a periodic input with waveform *S*_*e*,*i*_(*t*) *= S* sin(2 p*ω*_*stim*_*t*), where *S* is defined as the stimulation amplitude and *ω*_*stim*_ is the stimulation frequency. No stimulation is present for sub-cortical populations since non-invasive brain stimulation predominantly targets cortex.

For minimal thalamic drive (i.e. *D*_*LGN*_ *=* 1 × 10^-4^), the network engages intense resting state alpha activity at a frequency of about 8Hz due to the combined action of slow spike-frequency dependent adaptation and finite conduction velocity – resulting in propagation delays enhancing the prevalence of correlated rhythmic states (Lefebvre et al. 2011, Deco et al. 2009). The stability of the resulting oscillatory solutions is maintained and amplified due to thalamo-cortical feedback, leading to phase locked dynamics across all neural populations involved, both cortical and sub-cortical. The activity of each population in the resting state is depicted in Figure 1B. Parameter values chosen were aligned within the physiological range in the literature and are summarized in Table 1. The response of the sub-cortical populations (i.e. LGN, RTN) can be seen to adopt a stable phase offset with respect to cortical activity. Such phase differences have been observed experimentally and shown to be sensitive to ongoing oscillatory state (Slézia et al. 2011). In our model, this phase lag occurs because of the propagation delay to and back from the cortex. When thalamic drive is increased (i.e. *D*_*LGN*_ *=* 1), global oscillations are suppressed through a noise-induced Hopf bifurcation (See below). Through this transition, noise triggers a gradual change in effective gain between populations suppressing global oscillations and replacing them by asynchronous activity.

**Table 1:** Model Parameter Definitions and Values.

| Symbol | Definition | Value |
| --- | --- | --- |
| $\Omega$ | Network spatial extent | 1 |
| $N_e$ | Number of excitatory cells | 800 |
| $N_i$ | Number of inhibitory cells | 200 |
| $N_{th}$ | Number of thalamic cells | 200 |
| $N_{rtn}$ | Number of reticular cells | 200 |
| $\beta$ | Neural response function gain | 150 |
| $h$ | Firing rate threshold | 0.1 |
| $f_o$ | Maximal firing rate | 0.2 |
| $\tau_m$ | Membrane time constant | 1 |
| $\alpha_e$ | Membrane rate constant – excitatory cortical cells | 0.9 |
| $\alpha_i$ | Membrane rate constant – inhibitory cortical cells | 1.3 |
| $\alpha_{th}$ | Membrane rate constant – thalamic neurons | 0.5 |
| $\alpha_{rtn}$ | Membrane rate constant – reticular cells | 0.5 |
| $a$ | Neural adaptation rate constant | 0.01 |
| $b$ | Neural adaptation gain | 0.3 |
| $I_e$ | Constant current bias | 0 |
| $I_i$ | Constant current bias | -0.3 |
| $I_{th}$ | Constant current bias | -0.3 |
| $I_{rtn}$ | Constant current bias | -0.3 |
| $\tau_{th}$ | Thalamo-cortical delay | 45ms |
| $\tau_{rtn}$ | Reticular-thalamic delay | 10ms |
| $c$ | Conduction velocity | 0.35 m/s |
| $\rho$ | Connection probability | 0.2 |
| $w_{e \rightarrow e}^o$ | Synaptic connection strength | 20.4 |
| $w_{e \rightarrow i}^o$ | Synaptic connection strength | 30.6 |
| $w_{i \rightarrow e}^o$ | Synaptic connection strength | -30.6 |
| $w_{i \rightarrow i}^o$ | Synaptic connection strength | 20.4 |
| $w_{e \rightarrow lgn}^o$ | Synaptic connection strength | 34 |
| $w_{e \rightarrow rtn}^o$ | Synaptic connection strength | 34 |
| $w_{lgn \rightarrow e}^o$ | Synaptic connection strength | 85 |
| $w_{lgn \rightarrow i}^o$ | Synaptic connection strength | 85 |
| $w_{lgn \rightarrow rtn}^o$ | Synaptic connection strength | 34 |
| $w_{rtn \rightarrow lgn}^o$ | Synaptic connection strength | -34 |
| $\sigma_{e \rightarrow e}^2$ | Synaptic connection range | 0.01 |
| $\sigma_{e \rightarrow i}^2$ | Synaptic connection range | 0.01 |
| $\sigma_{i \rightarrow e}^2$ | Synaptic connection range | 0.25 |
| $\sigma_{i \rightarrow i}^2$ | Synaptic connection range | 0.25 |
| $\sigma_{e \rightarrow th}^2$ | Synaptic connection range | 0.01 |
| $\sigma_{e \rightarrow rtn}^2$ | Synaptic connection range | 0.01 |
| $\sigma_{th \rightarrow e}^2$ | Synaptic connection range | 0.25 |
| $\sigma_{th \rightarrow i}^2$ | Synaptic connection range | 0.25 |
| $\sigma_{th \rightarrow rtn}^2$ | Synaptic connection range | 0.25 |
| $\sigma_{rtn \rightarrow th}^2$ | Synaptic connection range | 0.25 |
| dt | Integration time step | 0.1 |

### Simulated EEG

In our model, encephalographic (EEG) dynamics is modelled by a weighted sum over somatic excitatory and inhibitory potentials, that is

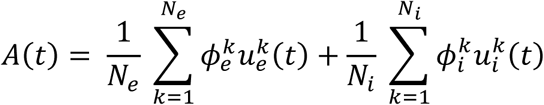

where 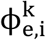 are real positive coefficients. Here we assume that the network fine scale structure is unknown, and thus consider random weights i.e. 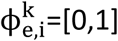. We, however, note that specific choices of the 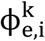 distributions can be made to increase the similarity of the network activity to physiological signals such as LFPs and EEG (Lindèn et al. 2010, Mazzoni et al. 2015).

### Spectral Analysis and Phase Distributions

Spectral analysis was performed using a fast Fourier transform(FFT) routine using freely available C++ scripts (Press et al, 2007). To quantify entrainment, we used a direct Fourier transform (DFT) with to compute the phase of the firing rate activity for cortical excitatory cells across independent trials, and relate this phase with the phase of the stimulation applied. As such, we computed the response of the network in rest and task conditions to stimulation delivered with a random phase for 200 independent trials of 2 seconds. In each trial, we randomly selected a time window of 500ms and computed the phase difference, at the stimulation frequency of 11Hz, between firing rate response and input stimuli. We then evaluated the distribution of these phase differences across all trials in both conditions. Results are shown in Figure 3E and 3F. In each trial, cortical neurons were driven with a 11Hz stimuli applied at a random phase.

### Mutual information

We computed mutual information between the stimulation signal *S*(*t*) and the firing rate activity to measure how well the stimulation waveform was reflected in the spiking patterns of cortical neurons. We assumed that for sufficiently high firing rates,the random fluctuations impacting the network responses can be approximated by Gaussian white noise, and computed,

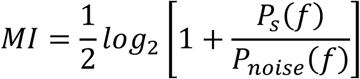

where ral density of the averaged network firing rates and deviover independent trials and the

### Noise-induced transition and resonance curves for a simplified delayed neural oscillator

The thalamo-cortical system has been thoroughly studied, both in experiments and models, and its low frequency dynamics have been shown to be largely determined by the presence of a delayed feedback loop between cortex and thalamus (Roberts & Robinson 2008). A thorough derivation of the reduced dynamics for the detailed spiking model we used in the simulations is far beyond the scope of this paper, and is highly challenging due to the combined presence of sparse and topographic synaptic projections, multiple cell types as well as spiking activity. To better understand the mechanism involved in shaping resting state cortical oscillations, we however built a simplified conceptual neural oscillator model whose limit cycle solutions emerge due to the combined presence of delayed feedback and slow spike-frequency adaptation,

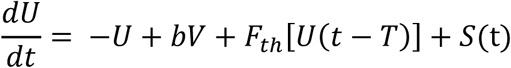

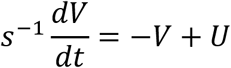

With feedback response function 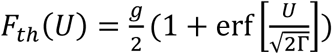 for some gain *g* < 0 and time delay *T*. The membrane potential proxy *U* denotes here the mean somatic membrane potential of cortical neurons, subjected to re-entrant inputs back from the thalamus. The response function also depends on the state-dependent noise variance Γ. This results from non-linear interactions between noise and a sigmoidal non-linearity, which is revealed by performing mean-field reduction (Lefebvre et al. 2015, Hutt et al. 2016). Note that a similar model has been used before to study state dependent entrainment and outlasting effects observed in experimental data in humans (Alagapan et al. 2016).

Assuming slow adaptation (*s* large), the smoothing of the neurons’ response functions (Lefebvre et al. 2015) leads to the effective linear dynamics with periodic forcing,

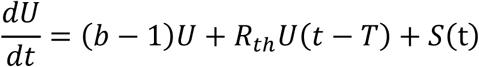

with 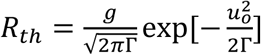 and fixed point *U*_0_. It has been shown that the value of the gain *R*_*th*_ is inversely proportional to the intensity of noise in the system. Indeed, for low values of Γ (i.e. in the rest state), |*R*_*th*_| remains high due to the steepness of the neuron response function. In contrast, as G inreases (i.e. in the task state), |*R*_*th*_| decreases (Hutt et al. 2016). The impact of state-dependent noise on oscillatory solutions can thus be quantified by analyzing how the stability of the system depends on the parameter Γ. It can be shown that increasing the noise variance in the system above causes a destabilization of limit cycle solutions through a supercritical Hopf bifurcation. Setting *S =* 0 and using the ansatz *u = u e*^-*λt*^ with *λ* = ±*i ω*_*c*_, where *ω*_*c*_ in the linearized system above, one obtains after separating real and imaginary parts,

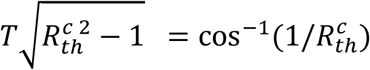

Setting *T =* 2 *τ*_*th*_ *=*90ms (i.e. twice the thalamo-cortical delay), one then obtains 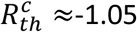.

The derivation of the resonance curve for a linear delayed system with periodic forcing is easily accomplished by finding the amplitudes of its solution, which can be computed explicitly via substitution. Let us assume an entrained oscillatory solution of the form

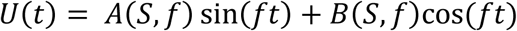

Substituting this ansatz in Eq. () above and solving for the *A*(*S*, *f*) and *B*(*S*, *f*), one can then compute the amplitude of the solution *u* as

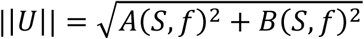

Where

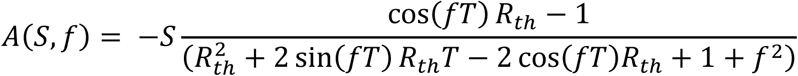

and

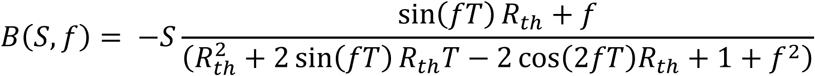

## ACKNOWLEDGEMENTS

This work has been supported by the Natural Sciences and Engineering Research Council of Canada (JL).

